# Balancing nutrient remobilization and photosynthesis: the dual role of lupin cotyledons after germination

**DOI:** 10.1101/2025.04.15.648908

**Authors:** Cecile Angermann, Björn Heinemann, Bianca Bueno Nogueira, Hans-Jörg Mai, Petra Bauer, Tatjana M. Hildebrandt

## Abstract

Efficient nutrient mobilization from seed storage tissues is essential for seedling establishment, particularly in legumes such as *Lupinus albus* (white lupin), which thrive in nutrient-poor soils. This study investigates the role of cotyledons in nitrogen (N) and mineral remobilization after germination during their transition from storage organs to photosynthetically active tissues including the metabolic challenges posed by coexistence of these two functions in epigeal germination. We cultivated white lupin seedlings under nitrogen-deficient conditions, analyzing cotyledon composition and function over 28 days. Our results indicate that 60 % of cotyledon-stored proteins are degraded within the first eight days, with free amino acids transiently accumulating before being redistributed to support growth. The progressive depletion of cotyledon reserves was accompanied by structural and metabolic changes, including an increase in photosynthetic proteins. However, cotyledon photosynthetic capacity remained lower than that of true leaves, suggesting a transient role in energy metabolism. The loss of cotyledons before day 12 significantly impaired seedling development, emphasizing their critical contribution to nitrogen, phosphate, and micronutrient supply during early growth. Comparative proteomic analysis revealed dynamic shifts in nutrient transport, amino acid metabolism, and stress response pathways following cotyledon removal. These findings underscore the significance of cotyledon nutrient remobilization in legume adaptation to low-fertility soils and highlight potential targets for breeding strategies aimed at improving nutrient use efficiency. By optimizing cotyledon nutrient composition and function, future breeding efforts could enhance seedling vigor, reduce fertilizer dependency, and improve the nutritional value of lupin-based foods.

**Significance statement:** This study uncovers how white lupin cotyledons balance nutrient storage and photosynthesis during early seedling development, ensuring sustained nitrogen and mineral supply under nutrient-deficient conditions. By detailing the proteome reorganization and consequences of premature cotyledon loss, it identifies key mechanisms supporting seedling resilience and adaptation to low-fertility soils.

## Introduction

Pulse legumes are vital for global nutritional food security and, as they are rich in protein and micronutrients, often considered as meat replacement. Especially increasing biodiversity through orphan legumes is important for the transition to sustainable agriculture, as these plants improve soil fertility through nitrogen fixation and thrive in diverse climates with low resource input. White lupin (*Lupinus albus*) is a proteinaceous and nutritious orphan crop in the lupin genus with relatively high protein and mineral contents, such as phosphate-containing compounds, iron (Fe), and zinc (Zn) (Karnpanit et al. 2017, Pereira et al. 2022, Spina et al. 2024). White lupin is particularly well adapted to nutrient-poor soils due to its specialized root system and efficient nutrient remobilization strategies. As legumes, lupins are able to acquire nitrogen from the environment through symbiotic fixation (Lucas et al. 2015). In addition, *Lupinus albus* can remobilize soil phosphates very efficiently by the formation of specialized cluster roots (Xu et al. 2020, Pueyo et al. 2021). However, seedling establishment still presents a significant challenge in nutrient-poor soils since plants rely on internal nutrient reserves before the specialized root system becomes fully functional. This dependency places specific demands on the quantity, composition, and metabolism of seed storage compounds, which have not been fully understood.

The cotyledons represent the primary nutrient store in lupin seeds. Upon seed maturation, storage proteins and lipids are deposited in protein storage vesicles and oil bodies within the cotyledon cells (Borek et al. 2009). In addition, *L. albus* seeds contain carbohydrates, mostly cellulose and oligosaccharides of the raffinose family, which are non-digestible in humans and other monogastric animals (Gdala and Buraczewska 1996, Sanyal et al. 2023). During germination, raffinose is hydrolyzed to sucrose and galactose, which can then serve as a source of energy and as a precursor for cellulose synthesis (Elango et al. 2022, Sanyal et al. 2023). The stored lipids representing 7 – 14 % of the seed dry weight are rapidly hydrolyzed by lipases and the free fatty acids are further metabolized to acetyl-CoA and NADH via β-oxidation in the peroxisomes (Borek et al. 2012, Baker et al. 2006, Graham 2008). The glyoxylate cycle converts acetyl-CoA to succinate for the synthesis of carbohydrates during gluconeogenesis and citrate can be exported from the peroxisomes into the mitochondria for ATP production (Baker et al. 2006, Pracharoenwattana et al. 2005, Pracharoenwattana and Smith 2008). White lupin seeds are remarkably rich in protein (30 – 40 %), yet, they do not primarily rely on amino acids from storage protein breakdown for energy production or anabolic processes during germination and early seedling establishment (Angermann et al. 2024). Instead, the total protein content remains relatively stable until the seedling emerges from the soil and thus the cotyledons can serve as a nitrogen store for the developing plant after the germination process has been completed. This delayed utilization of storage proteins suggests a highly regulated nutrient allocation strategy with a strong impact on seedling development on nutrient poor soils that warrants further investigation.

After initially acting as nutrient reserves cotyledons undergo a dynamic transformation during seedling development. Since lupins follow an epigeal germination strategy, their hypocotyl elongates causing the cotyledons to raise above the ground and become photosynthetically active before eventually senescing (Lovell et al. 1970). This transition to photosynthetic activity represents a major metabolic shift and allows cotyledons to contribute additional energy and carbon resources, reducing immediate dependence on stored metabolites. In contrast, hypogeal germination, characteristic of many other legumes, keeps cotyledons underground, where they function solely as storage organs before being fully depleted. Thus, white lupin presents an intriguing case where cotyledons assume a dual role, retaining large reserves of storage proteins and free amino acids while simultaneously becoming photosynthetically active. The coexistence of these two functions poses unique physiological and metabolic challenges, which require further investigation. How do lupin cotyledons regulate the shift between these roles without compromising either function? What molecular and enzymatic mechanisms orchestrate the gradual degradation of storage proteins while enabling chlorophyll accumulation and photosynthetic activation? Furthermore, how does the retention of storage compounds influence the efficiency of photosynthesis in cotyledons compared to true leaves? Understanding the mechanisms underlying this dual-function adaptation in lupin cotyledons could not only enhance our knowledge of seedling physiology but also inform breeding strategies aimed at improving crop resilience and nutrient efficiency.

Another aspect remaining understudied is the dynamics of mineral utilization from cotyledons to the developing plant in white lupin. Page et al. (2006) investigated the uptake and distribution of heavy metals in lupins during germination. However, their approach focused solely on uptake of some micronutrients and heavy metals via the roots and was not designed to assess the importance of the stored minerals for the transformation of cotyledons from storage into photosynthetic organ nor for the developing plants. This highlights the need for further research into mineral redistribution from cotyledons during germination and seedling establishment.

This study provides new insights into the dual function of *Lupinus albus* cotyledons, revealing how they balance nutrient storage and photosynthesis to support seedling establishment under nitrogen-deficient conditions. It explores the role of the cotyledons in nutrient supply beyond seedling establishment, with a focus on nitrogen allocation, mineral redistribution, and the impact of premature cotyledon loss. White lupin cotyledons retain a significant proportion of their stored proteins well beyond germination, suggesting a strategic allocation process in which major nitrogen stores are conserved ensuring sustained nutrient supply. Our research also dissects the dynamic reorganization of the cotyledon proteome during the transition from storage organ to photosynthetic tissue and explores how premature cotyledon loss triggers widespread metabolic and stress responses.

## Results

### Exploring the role of the cotyledons after germination

Previous research indicates that while storage lipids play a significant role during germination, storage proteins are primarily degraded at later developmental stages (Angermann et al. 2024). This suggests that proteins serve a different physiological purpose beyond germination. To explore this further, we cultivated *L. albus* for 28 days under nitrogen-limited conditions without rhizobia (which fix atmospheric nitrogen) and without nitrogen fertilization (Fig. 1, Supp. Fig. S1). This experimental setup ensures that the plants rely entirely on their seed reserves for nutrient supply. The cotyledons were removed and analyzed at five different timepoints (8, 12, 16, 20, 28 days after sowing, Fig. 1 A,B,D) and the phenotype as well as composition of the remaining plant were assessed at day 28 (Fig. 1 C,E).

**Fig. 1:**
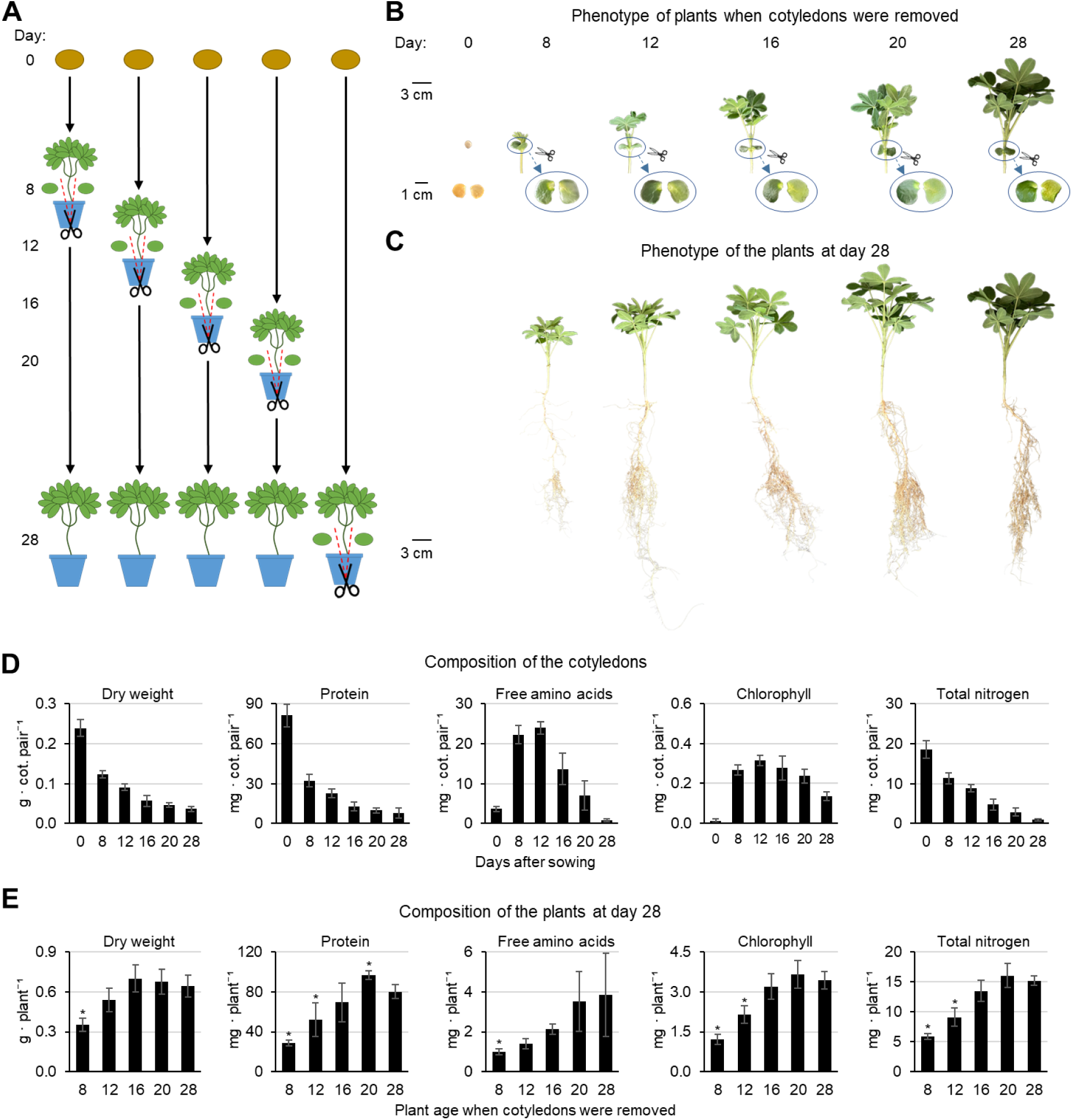
Exploring the role of the cotyledons after germination. (**A**) Scheme of the experimental set-up. Plants were grown for twenty-eight days and cotyledons were removed at day 8, 12, 16, 20 or 28 (control) after sowing. (**B**) Photos of shoots and cotyledons of representative plants at the time of cotyledon removal. Cotyledons: left: top view, right: bottom view. (**C**) Phenotype of representative plants at day 28 after sowing. (**D**) Biomass and composition of the cotyledons at day 0, 8, 12, 16, 20 and 28 after sowing. Data presented are means ± SD (n = 5). (**E**) Biomass and composition of the plants at day 28 after sowing. Data presented are means ± SD (n = 5). *significantly different to control (28d) (p<0.05). The complete dataset is provided as Supplementary dataset S1.

The cotyledons lost half of their total biomass and 60 % of their protein content already during the first eight days of germination and seedling establishment (Fig. 1D). High levels of free amino acids transiently accumulated with a peak in the cotyledons of juvenile plants between day 8 and 12. The total nitrogen content of the cotyledons continuously decreased over the 28 days and finally reached 5 % of the original seed content. Removing the cotyledons after day 12 did not lead to any significant changes in plant biomass, total nitrogen, protein, or amino acid content compared to the control (28d) (Fig. 1E). Cotyledon loss at day 8 resulted in a drastic reduction of the plant size as well as nitrogen metabolite contents at day 28 (Fig. 1 C,E). The total nitrogen content of the 4-week-old plant was 62 % lower than in the control (5.9 ± 0.5 vs. 15.2 ± 0.8 mg, Fig. 1E) and the missing fraction corresponded almost exactly to the nitrogen content of the removed cotyledons (11.3 ± 1.4 mg, Fig. 1D). The effect of cotyledon loss at day 12 was less pronounced but of a similar quality. Plants remained smaller with a lower protein and amino acid content than controls.

### Reorganization of the cotyledon proteome during transformation from storage organ to photosynthetic tissue

In order to monitor the transformation of the cotyledons from a storage organ to a photosynthetic tissue we analyzed changes in the proteome composition using our manually curated *Lupinus albus* proteome database for functional protein annotation (Angermann et al. 2024). The cotyledons represent a large part of the seedling at early developmental stages (65 % of the dry weight at day 8 and 42 % at day 12) but only a minor fraction in the mature plant (Fig. 2A). The seed proteome is dominated by storage proteins (28 %), LEA proteins (6 %) and proteins of the storage oil bodies (oleosins) (10 %) (Fig. 2C,D). Half of these major seed protein categories are degraded within the initial 8 days of development. 44 % of the nitrogen liberated by this proteolytic process is stored in the free amino acid pool mainly in the form of asparagine (Fig. 1D, Fig. 4A, Supplementary datasets S1 and S2) and the rest is directly reinvested into new functional proteins. The most pronounced reorganization of the cotyledon proteome occurs during the first 8 days of development (Fig. 2B,C,D). Enrichment analysis of functional annotations indicates a strong induction in primary metabolism including amino acid metabolism, cellular respiration, lipid degradation, nucleotide synthesis as well as photosynthesis and the production of the required cofactors (tetrapyrrole and isoprenoids, Fig. 2E) during seedling establishment. Subsequently, the relative composition of the proteome does not change as drastically (Fig. 2D). However, the protein fraction required for photosynthesis continuously increases during further development of the cotyledons until day 16 (Fig. 2D, Supp Fig. S2). The functional category “transport” is significantly enriched in all comparisons indicating dynamic changes in the transporter profile of the cotyledons throughout plant development. This includes an early up-regulation of proteins involved in mobilization and reallocation of iron (Fe) or regulation thereof, such as orthologs of nicotianamine synthase AtNAS1, transporters for Fe, nicotianamine-Fe and nicotianamine-Cu, as well as a transporter for Fe signals (YSL6, NRAMP3, OPT3; Supplementary dataset S2). These findings suggest that Fe and metal ion homeostasis is also a relevant feature of cotyledon transformation. During cotyledon senescence proteins involved in nutrient signaling and transport as well as cell wall metabolism are increased (Fig. 2D, Supplementary dataset S2).

**Fig. 2:**
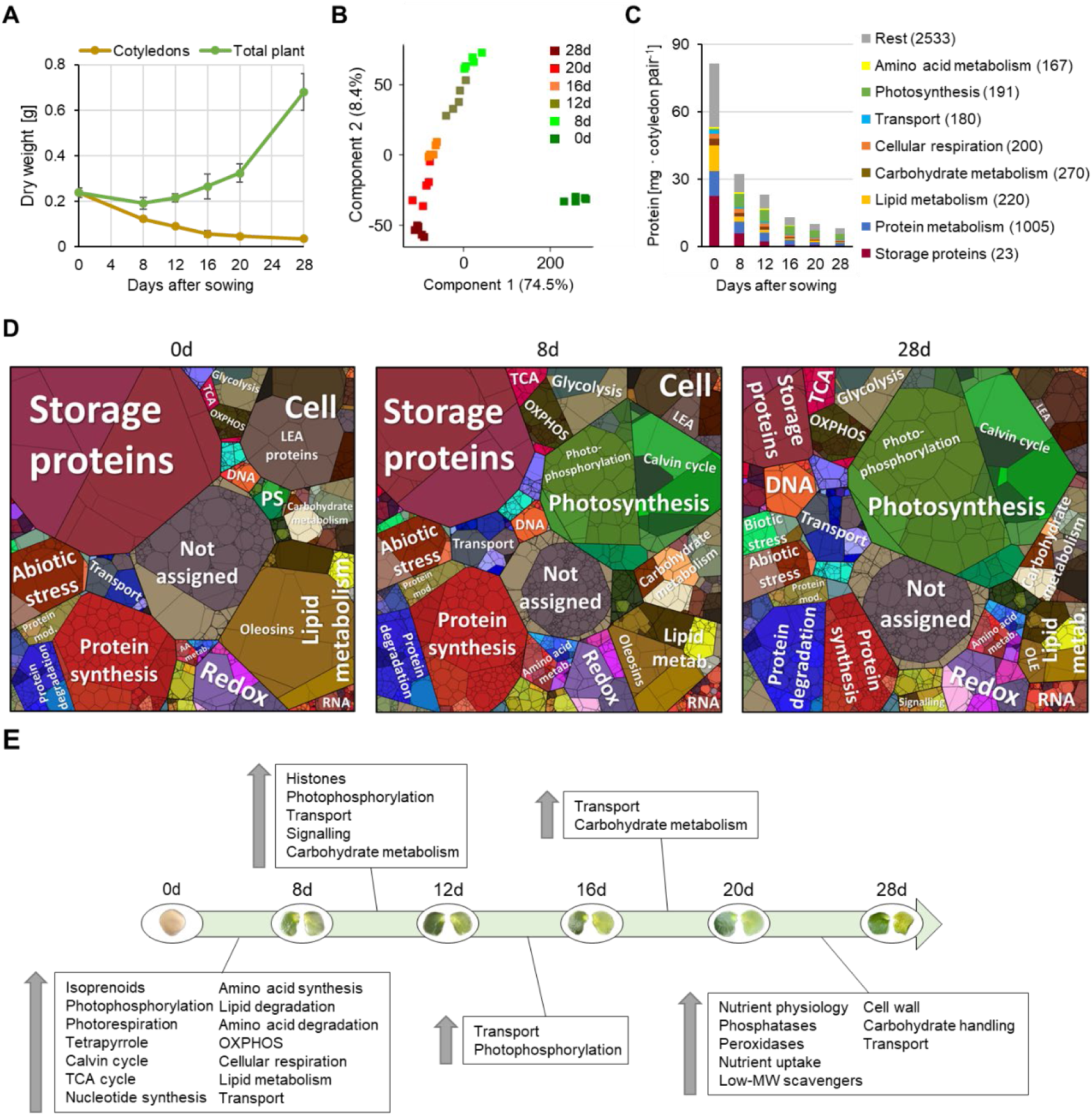
Reorganization of the cotyledon proteome during transformation from storage organ to photosynthetic tissue. (**A**) Biomass of lupin plants and cotyledons at 0 - 28 days after sowing. Data presented are means ± SD (n = 5). (**B**) Principal component analysis of the lupin cotyledon proteomics dataset. Data presented are means of five biological replicates. (**C**) Total protein content of the cotyledons and mass fractions covered by the major functional categories at day 0 - 28 after sowing. Numbers in parentheses indicate the number of different protein groups detected in the respective category. Mass fractions (protein abundances obtained by MS [iBAQ], multiplied with protein molecular weight) for all individual proteins are provided in Supplementary dataset S2. Data presented are means of five replicates. (**D**) Proteomaps illustrating the quantitative composition of the cotyledon proteome at day 0, 8, and 28 after sowing. LEA – LEA proteins; metab. – metabolism; mod. – modification; OLE – oleosins; PS - Photosynthesis. Proteins are shown as polygons whose sizes represent their mass fractions (see C). Proteins involved in similar cellular functions are arranged in adjacent locations and visualized by colors. Proteomaps were produced using the tool provided at https://bionic-vis.biologie.uni-greifswald.de/ (Liebermeister et al. 2014). Data presented are means of five biological replicates. (**E**) Significant changes in the cotyledon proteome during development from storage organ to photosynthetic tissue. Relative changes in abundance between subsequent sampling timepoints were analyzed for each individual protein group. Enrichment of functional annotations in the subset of proteins with significantly increased abundance compared to the total proteome was analyzed for each comparison, and significantly enriched categories are listed in the boxes in the order of descending enrichment factors. Data presented are means of five biological replicates. The complete dataset is provided as Supplementary dataset S2.

A direct comparison of the cotyledons to the true leaves at day 12 after sowing shows that while the leaves have an even higher protein content than the cotyledons at this stage, the cotyledons contain almost twice as much free amino acids (Fig. 3A, Supp. Fig. S3). The proteome composition is clearly different in these two tissues and shows that although photosynthetically active the cotyledon metabolism is dominated by catabolic pathways such as lipid, protein, and amino acid degradation as well as mitochondrial respiration (TCA cycle and OXPHOS, Fig. 3C,D). Proteins related to photosynthesis represent 22 % of the total protein mass in the cotyledons compared to 40 % in the leaves, which also contain a 4-fold higher chlorophyll content combined with a larger area (8.0 ± 1.5 vs. 3.0 ± 0.3 cm^2^) than the cotyledons (Fig. 3A). Comparison of photosynthetic performance using the light saturation curve of photosynthesis shows much higher electron transport rates in leaves at all light intensities (Fig. 3B). Comparable results were obtained by comparing the net photosynthetic rates of leaves and cotyledons. At a photon flux density of 500 µmol photons · m^-2^ · s^-1^, net photosynthetic oxygen production in leaves is 9 times higher than in cotyledons (23.0 ± 4.5 vs. 2.6 ± 1.8 µmol oxygen · min^-1^ · g^-1^ DW) (Supplementary dataset S3). Interestingly, the maximum quantum yield of photosystem II (cotyledon: 0.846 ± 0.005; leaf: 0.848 ± 0.004) is similar in both tissues at 12 days after sowing (Supplementary dataset S3). In addition to photosynthesis the sulfur assimilation pathway as well as protein, lipid and amino acid synthesis reactions are enriched in the leaves compared to the cotyledons (Fig. 3 C,D).

**Fig. 3:**
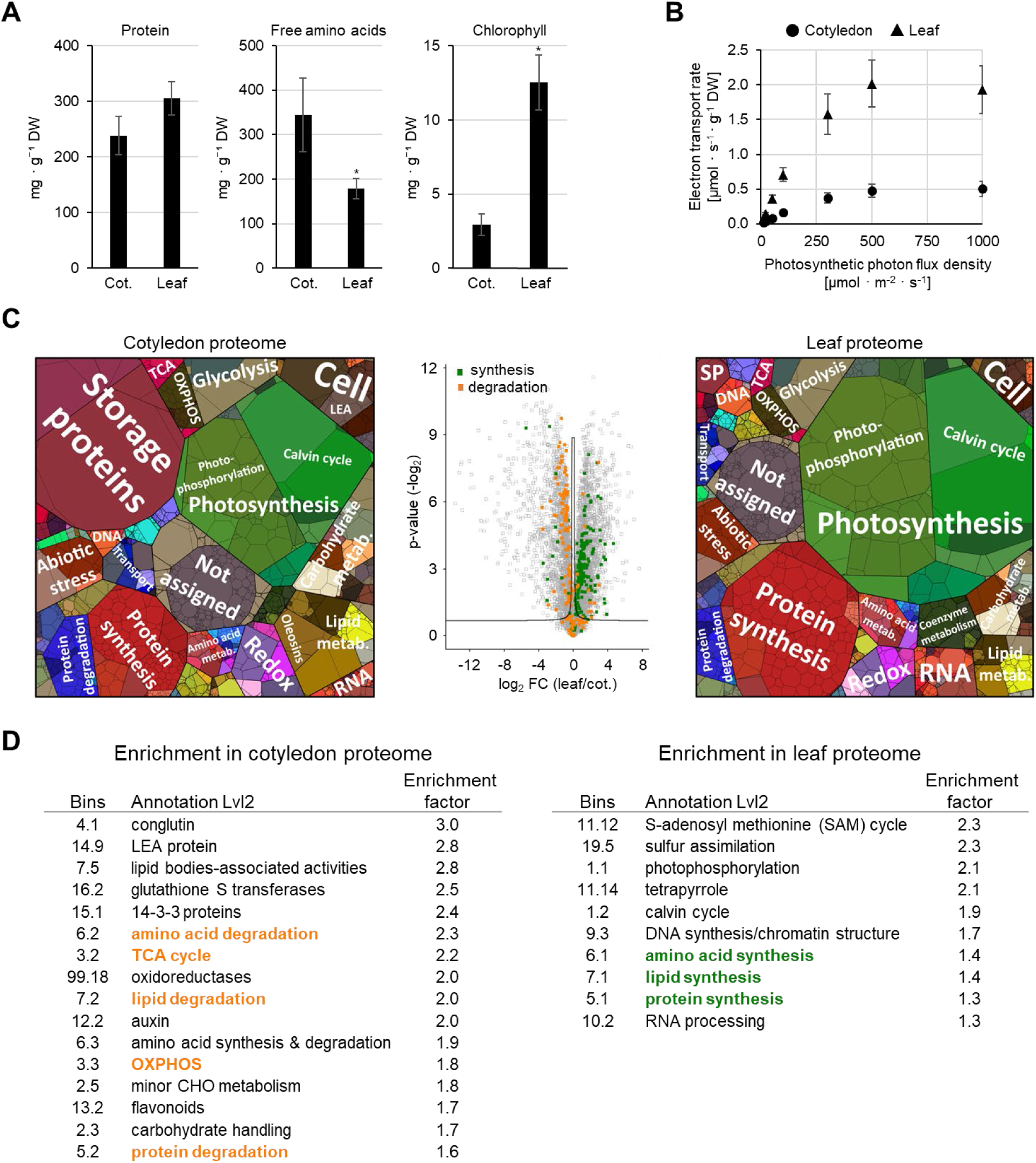
Proteome composition and photosynthetic performance of cotyledons vs. true leaves in a lupin plant at 12 days after sowing. (**A**) Biomass and composition of the cotyledons and the true leaves. Data presented are means ± SD (n = 5). *significantly different to control (cotyledon) (p<0.05). (**B**) Light saturation curve of photosynthesis. Data presented are means of five biological replicates. (**C**) Comparison of the quantitative and relative composition of the cotyledon proteome and the leaf proteome. LEA – LEA proteins; metab. – metabolism; SP – Storage proteins. Proteomaps illustrate the quantitative proteome composition. Proteins are shown as polygons whose sizes represent their mass fractions. Proteins involved in similar cellular functions are arranged in adjacent locations and visualized by colors. Proteomaps were produced using the tool provided at https://bionic-vis.biologie.uni-greifswald.de/ (Liebermeister et al. 2014). The volcano plot illustrates log2 fold changes (FC) in relative protein abundances between the two tissues. The significance threshold is indicated by solid lines. Proteins of catabolic pathways enriched in the cotyledons are highlighted in orange and proteins of synthesis pathways enriched in the true leaves are highlighted in green (see D). Data presented are means of five biological replicates. (**D**) Enrichment of functional categories in proteins that are significantly increased in the cotyledon proteome (left) or the leaf proteome (right) at day 12 after sowing. Data presented are means of five biological replicates. The complete dataset is provided as Supplementary dataset S3.

**Fig. 4:**
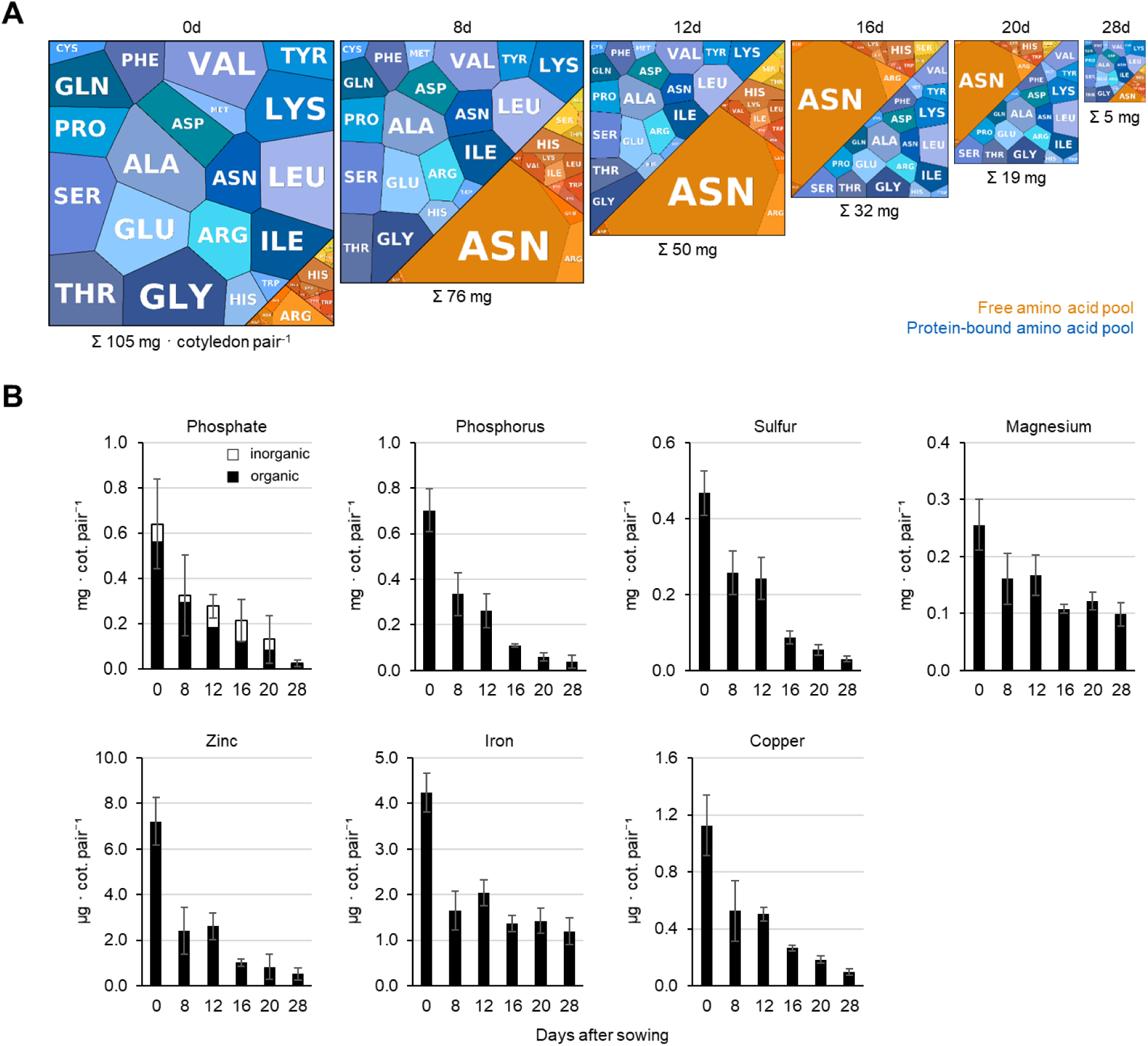
Remobilization of nutrients from the cotyledons. (**A**) Quantitative composition of the free (orange) and protein-bound (blue) amino acid pools in the cotyledons at day 0, 8, 12, 16, 20, and 28 after sowing. Amino acids are shown as polygons whose sizes represent the molar fractions. Free amino acid contents were quantified by HPLC, and the quantitative amino acid composition of the proteome was calculated on the basis of molar composition of the proteome (see Supplementary datasets S1 and S2). The area of the square corresponds to the total amino acid content per cotyledon pair at a given timepoint. Aminomaps were produced using the tool provided at https://bionic-vis.biologie.uni-greifswald.de/ (Liebermeister et al. 2014). The complete amino acid profiles are provided in Supplementary dataset S3. Data presented are means of five biological replicates. (**B**) Mineral composition of the cotyledons at day 0, 8, 12, 16, 20 and 28 after sowing. Data presented are means ± SD (n = 3 - 9). The complete dataset is provided as Supplementary dataset S1.

### Remobilization of nutrients from the cotyledons

Taken together the proteome datasets and amino acid profiles reveal that nitrogen remobilization from the cotyledons to the growing lupin plant proceeds in two stages. A large fraction of the protein store from the seed (60 %) is degraded already during germination and early seedling establishment (Fig. 1D, 2C). However, high levels of free amino acids in the cotyledons of young plants (> 200 mM at day 8) indicate transient storage of nitrogen in the free amino acid pool for later utilization in the growing plant (Fig. 1D, 4A). The exceptionally high share of free amino acids in the total pool persists in the cotyledons until day 20, whereas the total biomass and thus also the total amino acid content decreases drastically (Fig. 1D, 4A).

*L. albus* is well adapted to growing on soils with a low phosphate content due to its ability to develop specialized cluster roots (Xu et al. 2020, Pueyo et al. 2021). Thus, phosphate storage and supply to the young plant might also be an important function of the cotyledons. The phosphate store of the white lupin seeds analyzed in this study mainly consisted of organic compounds (Fig. 4B, phosphate). The total phosphate content of the cotyledons decreased continuously during the first four weeks of plant growth with the strongest drop (50 %) occurring during germination and seedling establishment until day 8. Cotyledon removal after day 12 had no significant effect on the total phosphate content of the plants at day 28. In contrast, loss of cotyledons at day 8 or 12 resulted in a drastic reduction in the plant’s phosphate content at day 28. The missing fraction corresponded to the phosphate content of the removed cotyledons (Supplementary dataset S1).

There are further important macro- and micronutrients present in the cotyledons that can serve in photosynthesis. To test whether they were utilized with a similar or different pattern from N-storage compounds and phosphate, we conducted an elemental analysis using the same approach as described above (Fig. 4B). As expected, the phosphorous (P) pattern followed that of phosphate and N. There was a continuous usage of P from the cotyledons during the 28 days, while no extra P was acquired from soil (Fig. 4B, phosphorus). Since phosphate is the P form in living organisms, this was expected. Surprisingly, the other tested elements showed evidence of two new nutrient usage patterns. Iron (Fe) and sulfur (S) are needed for electron transport in chloroplasts and energy production in mitochondria. Fe is also required as cofactor for chlorophyll biosynthesis, S and N assimilation. Both, Fe and S were nearly fully used from cotyledons during the first 8 days of germination prior to onset of photosynthesis in cotyledons (Fig. 4B, sulfur, iron). Already during this time frame, seedlings began acquiring Fe and S from the root substrate. In contrast, micronutrients like copper (Cu), zinc (Zn) and macronutrient magnesium (Mg) were used from cotyledons during the first 8 days, while only after that the increase in total plant contents must be through root uptake (Fig. 4B, copper, zinc, magnesium). Hence, there are different dynamics of mineral mobilization from cotyledons and soil, with at least three different patterns for mineral utilization from cotyledons over the 28-day time span.

Premature cotyledon loss had no significant effect on the relative mineral contents of the plants at day 28 (Supp. Fig. S4). However, since early removal of the cotyledons led to a strong growth retardation, the total mineral contents of the plants at day 28 were decreased by the same factor as the total biomass (Fig. 1E, Supplementary dataset S1).

### Responses of the *Lupinus albus* proteome to premature cotyledon loss

The results of this study indicate that the cotyledons are able to support plant growth by providing nutrients for about two weeks after germination in *L. albus* plants grown without external nutrient supply under control conditions. Loss of the cotyledons at day 16 or later did not lead to any significant changes in the plant phenotype (Fig. 1) or composition of the proteome (Fig. 5A) at day 28. In contrast, not only the growth phenotype (Fig. 1) and nutrient content (Fig. 1E, Fig. 4B) but also the proteome of plants grown without cotyledons from day 8 or 12 was clearly different from that of control plants with 1214 and 943 proteins of significantly changed relative abundance, respectively (Fig. 5A,B, Supplementary dataset S4). The profile of protein induction or repression was highly consistent and the same trend was also visible after later cotyledon loss (red and blue squares in Fig. 5A). Enrichment analysis indicates a relative increase in protein and lipid catabolism as well as nutrient signalling and transport processes in plants without cotyledons (Fig. 5B, left). The strongest enrichment was detected in proteins required for phosphate assimilation, highlighting the function of the cotyledons in phosphate storage. There was also an induction of certain metal homeostasis-related proteins, which may follow the rapid early translocation of metals from the cotyledons, accompanied by adjustments to an early onset of Fe uptake from soil (Supplementary dataset S4). In addition, a biotic stress response was induced on the proteome level. Surprisingly, removing the cotyledons also led to a higher relative content of storage proteins in the remaining plant tissues at the age of four weeks. A significant decrease was detectable for proteins related to photosynthesis and the production of the required cofactors, amino acid and protein synthesis, as well as carbohydrate and coenzyme metabolism (Fig. 5B, right). These changes in relative proteome composition provide an impression of which pathways are particularly relevant for adjusting to early loss of the cotyledons. However, taking total protein abundance into account shows that photosynthesis and protein metabolism quantitatively contribute by far the largest share to reducing the protein content of the plant after early cotyledon loss (Fig. 5C). The plant is able to save 15.5 mg protein by reducing its relative protein content by 35 % from 126 ± 16 to 82 ± 13 mg · g^-1^ DW after premature cotyledon loss at day 8 (Supp. Fig. S4, Supplementary dataset S4). The most effective nitrogen saving strategy, however, is the reduction in plant biomass from 644 ± 84 to 353 ± 47 mg dry weight that, based on the mean tissue protein content of the control plant, would save 36.7 mg protein.

**Fig. 5:**
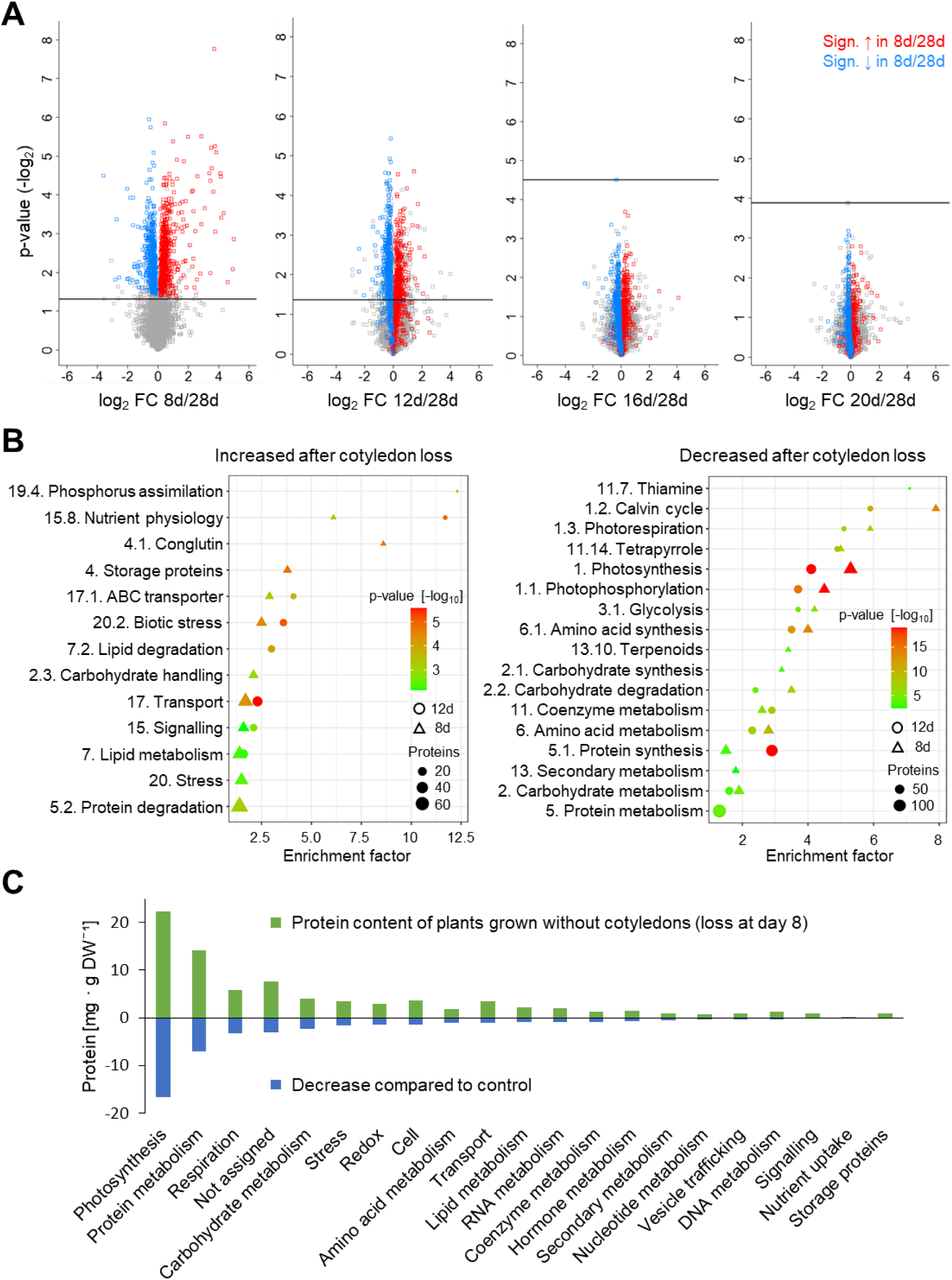
Response of the *Lupinus albus* proteome to premature cotyledon loss. (**A**) Volcano plots illustrating differences in the proteome of 4-week-old lupin plants grown without cotyledons from day 8, 12, 16, or 20 compared to control plants. Representative pictures of these plants are shown in Fig. 1C. To visualize the level of consistency in the effect of cotyledon loss on the plant proteome, proteins that are significantly increased or decreased after removal of the cotyledons at day 8 compared to the control are highlighted in red and blue, respectively, in all plots. The horizontal lines mark the significance threshold (p < 0.05, FDR: 0.1). Data presented are based on five biological replicates. (**B**) Enrichment of functional categories in proteins that are significantly increased (left) or decreased (right) in the plant proteome at day 28 after cotyledon loss at day 8 (triangles) or day 12 (circles). Data presented are based on five biological replicates. (**C**) Protein content of lupin plants at day 28 after removal of the cotyledons at day 8 (green bars) and difference to control plants (blue bars) for the major functional categories. All plants were harvested at day 28. Data presented are means of five biological replicates. The complete dataset is provided as Supplementary dataset S4.

## Discussion

The present study focuses on the function of the cotyledons in nutrient supply *Lupinus albus* after initial seedling establishment including their contribution to photosynthesis as well as nitrogen and mineral nutrition (Fig. 6). We investigate the timeframe of nutrient remobilization without external nutrient supply mimicking cultivation on nutrient-poor soils and monitor the integration of metabolic pathways with physiological requirements during cotyledon development from seed storage tissue to photosynthetic organ up to senescence. Our results reveal that *Lupinus albus* cotyledons provide essential nutrients during a critical phase of about two weeks after germination, with premature loss at day 8 or 12 causing significant changes in growth, nutrient content, and proteome composition, while later cotyledon removal has minimal effects on plant development.

**Fig. 6:**
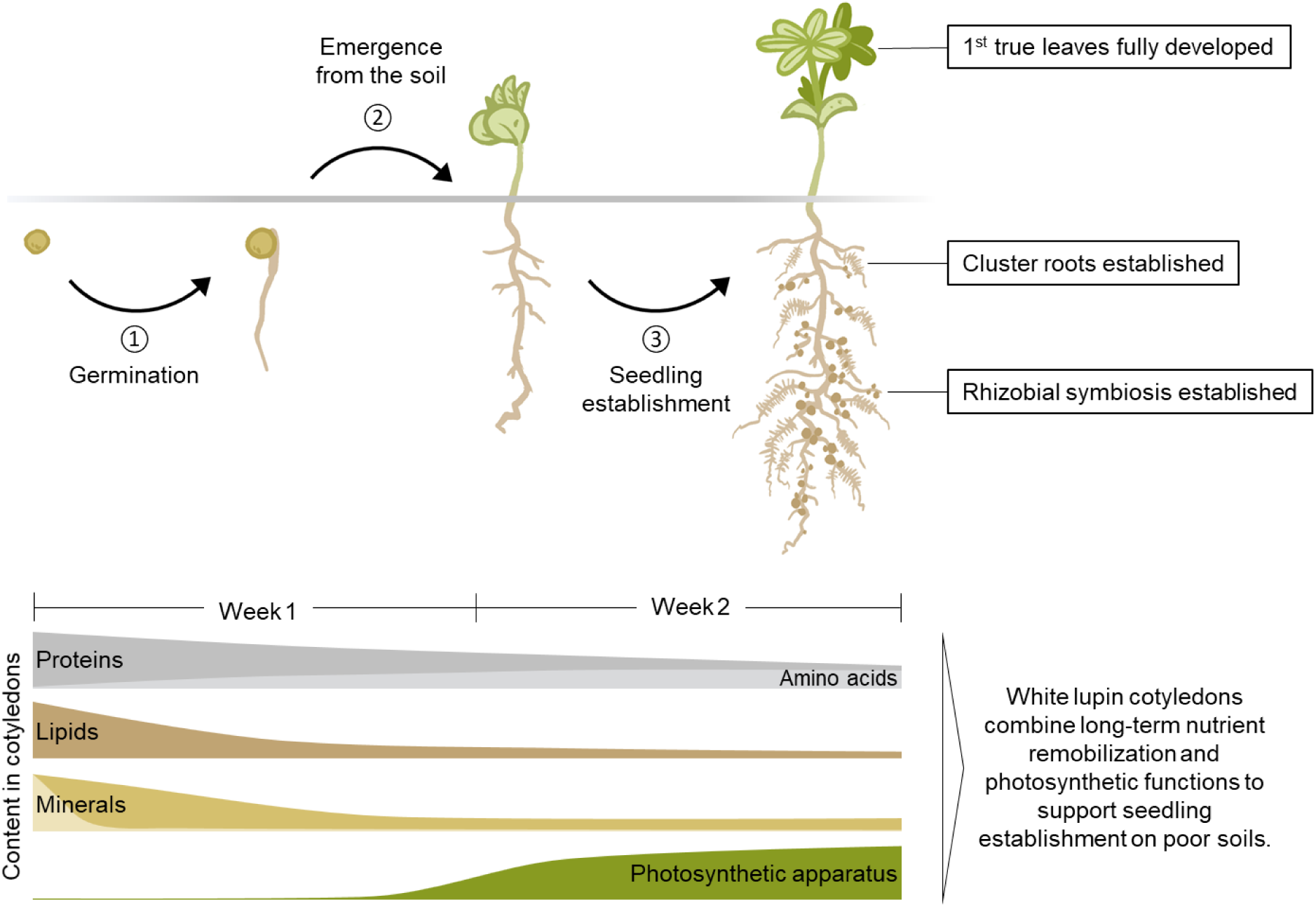
Combining nutrient remobilization and photosynthesis: the dual role of white lupin cotyledons during seedling establishment on nutrient-poor soils. Seed storage compounds provide energy and precursors for the synthesis of macromolecular structures during germination (1) and fully support plant growth until the seedling has emerged from the soil (2) and photosynthesis has been established. In some species including lupins the cotyledons are transformed into photosynthetically active leaves after germination allowing them to function as storage tissues for an extended period. The delayed remobilization of specific storage compounds, such as proteins and phosphate, is essential for successful seedling establishment (3) on nutrient deficient soils. The internal nutrient supply supports growth until specialized root structures and symbiotic associations are developed, enabling the seedling to become fully self-sufficient. In natural environments this is usually achieved within two weeks after germination (James et al. 1997, Neumann et al. 1999). Details on remobilization patterns of the different storage compounds and reorganization of the cotyledon proteome are provided in the main text.

### Cotyledons combine nutrient storage and photosynthetic functions in the young lupin plant

The cotyledons of *Lupinus albus* serve a dual role, initially functioning as storage tissue to fuel germination before emerging from the soil and becoming photosynthetically active. While supporting energy production, they also retain substantial reserves of storage proteins and free amino acids. This transformation requires a precise balance of nutrient remobilization and energy generation, which we analyzed through proteomics, metabolic, physiological, and functional studies over 28 days. Proteins related to photosynthesis are strongly induced already during the first days of seedling establishment, and the cotyledons become photosynthetically active and thus contribute to photoautotrophic ATP production and carbon fixation. However, compared to the true leaves they have a significantly lower photosynthetic capacity, although the efficiency of photosystem II is comparable. The proteome composition of cotyledons reveals a clear metabolic focus on catabolic processes and resource remobilization. Therefore, as soon as the first true leaves are fully developed their net photosynthetic rate exceeds the contribution of the cotyledons and the composition of their proteome is consistent with their dominant role as a source tissue for photosynthate and biosynthetic processes. Thus, white lupin cotyledons can be classified as semi-photosynthetic storage tissues with a prolonged dual function in nutrient remobilization and photosynthesis. Research on the role of soybean cotyledons in seedling establishment based on growth analysis had also indicated that cotyledon photosynthesis is not sufficient to increase plant growth, but can provide the carbohydrates needed to compensate for respiratory losses during early seedling establishment (Harris et al. 1986). A more recent study on castor bean cotyledons came to a similar conclusion demonstrating that cotyledon photosynthesis generated sufficient carbohydrates and energy to support the emergence of the first true leaf and sustained seedling growth until the leaf had fully expanded (Zheng et al. 2011). These examples indicate that cotyledon photosynthesis has a transient function during epigeal germination in species storing proteins as well as lipids.

### Long-term nutrient remobilization from the cotyledons to the lupin seedling enables growth on nutrient poor soils

A major storage compound in the lupin seed is nitrogen, mainly in the form of proteins and amino acids (82 % of total N). However, *Lupinus albus* does not use amino acids as a primary substrate for energy metabolism during germination (Angermann et al. 2024). Nitrogen remobilization from cotyledons proceeds continuously throughout the first weeks of plant development and can be divided into two steps. Within the first eight days of germination and seedling establishment, it is clearly associated with proteolysis and a significant transfer of amino acids from the protein-bound form to the soluble pool. The protein content is halved and at the same time, the amount of free amino acids strongly increases. Proteolysis releases large amounts of free amino acids, which are used as precursors for new functional proteins and nitrogenous metabolites, or accumulate in the free pool. Asparagine strongly accumulates and serves as a suitable form for storage and transport of reduced nitrogen due to its favorable C/N ratio, minimal net charge and low reactivity under physiological conditions. The conversion of protein-bound nitrogen to the soluble form as asparagine is likely to play a key role in the remobilization and transport of nitrogen to growing tissues (Lea et al. 2007). The synthesis of asparagine involves the assimilation of free ammonium by the combined reactions of glutamine synthetase and asparagine synthetase (Lea and Flowden 1975). These enzymes show a high relative increase in cotyledon protein content during seedling establishment (Supplementary Dataset S2). Quantitative evaluation of the seed and plant composition revealed that the proteins stored in the seeds were required almost exclusively for the synthesis of new functional proteins during seedling development. Additional nitrogen containing compounds such as chlorophyll and nucleotides quantitatively require a minor input of nitrogen resources. These results suggest that nitrogen stored in lupin cotyledons plays a crucial role in several aspects. Degradation of seed-stored proteins releases nitrogen to provide precursors for successful early germination, but also to provide storable but transportable soluble forms of nitrogen for longer periods to enable establishment on nitrogen-poor soils until the symbiosis with N2-fixing rhizobia has been established, which is usually achieved within 2 - 3 weeks after germination (James et al. 1997).

Another important macronutrient involved in several biological processes is phosphorus (Raghothama 2005). Phosphate is required for nucleic acids, membrane lipids, ATP as well as diverse metabolic and regulatory processes. Low phosphate availability in many natural ecosystems especially under acidic conditions promoting inorganic complexation is a major constraint on crop production. Short-term deficiencies can be compensated by intracellular stores, but long-term phosphate starvation leads to stress responses such as an altered root architecture, an increase in the root/shoot ratio, and an induction of phosphate uptake and recycling pathways (Raghothama 2005). In P-limited environments lupin seedlings rely on the phosphate stored in the cotyledons until specialized, densely packed lateral root structures called cluster roots are fully developed. This process usually requires 12 - 14 days after imbibition (Neumann et al. 1999). The cluster roots can mobilize soil phosphates very efficiently by the secretion of organic acids and acid phosphatases and an induction of a high-affinity P transport system (Xu et al. 2020, Pueyo et al. 2021). Our results show a continuous loss of phosphate from the cotyledons throughout plant development and a deficit in phosphate content in 28-day-old plants after early cotyledon loss. Induction of phosphate assimilation including phosphate transporters and regulatory proteins was the process showing the strongest enrichment in the lupin proteome in response to cotyledon removal within the first two weeks of growth. These results demonstrate that continuous phosphate supply from the cotyledons is able to bridge the time gap between germination and cluster root establishment and is thus essential for successful seedling establishment on soils with a low phosphate availability.

The remobilization of micronutrients, particularly iron (Fe), copper (Cu), and zinc (Zn), from cotyledons begins before nitrogen depletion in plants. Early metal translocation from cotyledons and Fe uptake from soil coincide with the activation of metal homeostasis proteins. In Arabidopsis thaliana, various transporters like AtYSL6 (Conte et al. 2013), AtMfl1 (Tarantino et al. 2012), AtNRAMP3 (Lanquar et al. 2005, Thomine et al. 2003, Li et al. 2018), and AtOPT3 (Khan et al. 2018) play crucial roles in metal distribution and mobilization. Early induction of Fe uptake supports critical processes like chlorophyll biosynthesis and seedling development, as Fe-containing enzymes are essential. The process ensures Fe retention in cotyledons and supports nitrogen assimilation and metabolic functions, highlighting its importance during and after germination.

### Premature cotyledon loss leads to nutrient deficiency and growth retardation

Premature cotyledon loss triggers profound phenotypic and molecular responses, with severity increasing the earlier the loss occurs. A major consequence is a significant reduction in plant growth, accompanied by a decrease in relative protein content in the remaining plant tissue by day 28. Nutrient deficiencies, particularly phosphate limitation in combination with nitrogen and iron scarcity, are known to reduce biomass accumulation (Pueyo et al. 2021). A loss of the cotyledons before cluster roots and the symbiosis with N2-fixing rhizobia have been established elicits severe growth retardation and specific proteomic adaptations to nitrogen and phosphate starvation. These adaptations include a relative decline in highly abundant pathways, such as photosynthesis and protein synthesis, both of which require substantial nitrogen investment and thus bind a large share of the available nitrogen resources. Conversely, proteins involved in resource management, including signaling and transport, show a relative increase, indicating a shift towards metabolic prioritization under nutrient constraints.

Previous studies have demonstrated that cotyledon loss negatively impacts early seedling establishment and can extend its effects into later developmental stages, leading to delayed flowering, reduced inflorescence production, and lower seed yield across various plant species (Hanley and Fegan 2007, Zhang et al. 2011, Hanley and May 2006, Wang et al. 2019). In *Arabidopsis thaliana*, some of these effects were partially mitigated by exogenous sucrose and auxin application (Wang et al. 2019). However, the extent of susceptibility to early cotyledon loss varies between species, suggesting that ontogenetic differences in seedling tolerance to tissue loss could significantly influence plant fitness in mature plant communities (Hanley and Fegan 2007). Seed size also plays a crucial role in tolerance; seedlings from larger seeds, such as white lupin, rely more heavily on their initial nutrient reserves and are therefore more sensitive to cotyledon loss than those from smaller seeds (Hu et al. 2017).

Tolerance and an efficient response to premature cotyledon loss can become physiologically relevant during herbivore attack. The lupin leaf weevils *Sitona gressorius* and *Sitona griseus* (Coleoptera: Curculionidae) cause considerable damage to lupin crops in Europe (Piedra-García and Struck 2021). In the spring, adult weevils start feeding on newly emerged plants and can already at this stage severely weaken the young seedlings by damaging the cotyledons (Ströcker et al. 2013). Subsequently, the soil-dwelling larvae penetrate and destroy the root nodules as well as the surrounding root tissue. In addition to the negative impact on symbiotic nitrogen fixation capacity the injuries caused by the feeding activities increase the risk of fungal infections (Piedra-García and Struck 2021). After removing the cotyledons, we consistently detected an induction of proteins associated with pathogen responses in the four-week-old lupin plants. Activation of plant innate immunity in response to wounding has mainly been studied in model species (Savatin et al. 2014). Damage-associated molecular patterns (DAMPs) elicit local and systemic responses mediated by a combination of electrical signals, calcium spikes, reactive oxygen species, and hormones (Farmer et al. 2020). Mechanical injury activates defenses that are similar to those induced by herbivores and lead to increased pathogen resistance (Orlovkis and Reymond 2000, Chassot et al. 2007). Our dataset demonstrates a long-lasting effect of mechanic cotyledon removal on the *L. albus* proteome. It needs to be elucidated whether this response protects the plants from weevil larvae and/or microbial pathogens and also whether it is specific for the cotyledons.

## Conclusions

In this study, we observed different roles of the cotyledon at different times of plant development. Initially, cotyledons act as storage tissues for a variety of nutrients. Catabolic processes, transport and the initialization of synthesis pathways are in the focus. Later, during epigeal germination, cotyledons act not only as nutrient stores but also as photosynthetic tissues, which contribute energy and carbohydrates for plant development and growth. Once the first true leaves have emerged, the photosynthetic aspects recede into the background and cotyledon senescence with catabolic processes and nutrient export take priority. Seed nutrient reserves are required for adequate plant growth and development, not only during germination or until the seedling emerges from the soil and becomes autotrophic through photosynthesis. The observed early Fe redistribution and protein analysis suggest that this transition is initiated very early after germination in the cotyledons. Internal nutrient supply is required for a prolonged period until specialized root structures and symbiotic interactions are established and the seedling becomes fully self-sufficient, even on nutrient-poor soils. Our results demonstrate the importance of the cotyledons of *Lupinus albus* during the first two weeks of seedling establishment. At this stage, premature cotyledon loss causes drastic phenotypic and molecular losses and responses, as well as activation of starvation and stress response pathways. The large nutrient stores in the seeds of *Lupinus albus* are thus prerequisite for thriving in nutrient-deprived environments.

Given the central role of legumes in sustainable food systems, unraveling the dynamics of cotyledon nutrient remobilization has significant implications beyond basic plant physiology. Optimizing this process could lead to the development of legume cultivars with improved seedling vigor, higher nitrogen use efficiency, and enhanced protein content, traits that are crucial for both agricultural sustainability and human nutrition. Breeding efforts focused on vigorous seeds with an optimal nutrient composition could improve not only seedling establishment but also the nutritional value of legume-based foods, addressing global challenges related to food security and dietary health. Additionally, understanding how cotyledon transformation contributes to early photosynthesis could offer novel strategies for enhancing seedling growth under suboptimal soil conditions. Beyond this, it would be thrilling to unravel the regulatory switches that determine whether a legume species uses epigeal or hypogeal germination patterns.

## Materials and Methods

### Plant material and growth conditions

White lupin (*Lupinus albus* cv. "Nelly") seeds were stored in the dark at 4 °C until use. Prior to cultivation, seeds were soaked in demineralized water at 20 °C for 16 h and then transferred to water-saturated expanded clay substrate (LamstedtDan, 4 - 8 mm, Fibo ExClay Deutschland GmbH, Lamstedt, Germany). Plants were grown in a phytochamber (22 - 24 °C, 16 h light, 8 h dark, light: 110 µmol photons · s^-1^ · m^-2^). On day 8, 12, 16 or 20 after sowing, the cotyledons were separated and harvested. The control group of plants retained their cotyledons until day 28. On day 28 after sowing, all plants were harvested. Except for cotyledon vs. leaf experiments, where the respective tissues were harvested at day 12 after sowing. At harvest, the remaining plant organs were frozen in liquid nitrogen. The plant material was lyophilized in an Alpha 1-2 LD+ freeze dryer (Christ, Osterode, Germany). The dried material was then ground into powder and used for the experiments. For one sample, several plants were pooled (10 seeds; 6 pairs of cotyledons at day 8; 5 pairs of cotyledons at days 12, 16 and 20; 4 pairs of cotyledons at day 28; 3 plants at day 28 with cotyledons separated at day 8, 2 plants each at day 28 with cotyledons separated at days 12, 16, 20 and 28). Five different pools were analyzed at each time point as biological replicates.

Plants for elemental analysis were grown as described above. Plant material was harvested at the same time points but dried at 60 °C for 2 days and ground to a fine powder.

Samples for analysis of photosynthetic parameters were grown as described above and used fresh immediately after harvest on day 12.

### Quantification of total carbon and nitrogen

The total carbon and nitrogen content of the samples was measured according to Andrino et al. (2019), using an Elementar vario MICRO cube C/N analyzer (Elementar GmbH, Hanau, Germany).

### Quantification of chlorophyll

The quantification of chlorophyll was carried out according to a modified version of the method described by Lichtenthaler (1987). 5 mg plant powder was dissolved in 700 ml methanol (100 %) and incubated for 20 min at 80 °C with shaking. After centrifugation (10 min, 4 °C, 18,800 xg), the chlorophyll content of the supernatant was quantified by plate reader (Multiskan Sky, Thermo Fisher Scientific, Dreieich, Germany) (wavelengths: 470 nm, 653 nm and 666 nm).

### Quantification of proteins

Proteins were extracted as described by Angermann et al. (2024). The Pierce BCA Protein Assay Kit (Thermo Fisher Scientific, Rockford, Illinois, USA) was used for protein quantification with globulin as a standard.

### Quantification of free amino acids

Free amino acids were quantified according to Batista-Silva et al. (2019). The pre-column derivatisation with ophthaldialdehyde (OPA) and fluorenylmethoxycarbonyl (FMOC) was based on the Agilent application note ’Automated amino acids analysis using an Agilent Poroshell HPH-C18 Column’. Samples were injected onto a Agilent 100 mm, 3 mm InfinityLab Poroshell HPH-C18 column (2.7 mm) (Agilent Technologies, Waldbronn, Germany) using an Ultimate 3000 HPLC system (Thermo Fisher Scientific, Germering, Germany) for cotyledon timeline experiments and the cotyledon removal experiment. Samples for the comparison of cotyledons with true leaves were analysed by an Agilent 1260 Infinity II HPLC system (Agilent Technologies, Waldbronn, Germany).

### Elemental analysis by inductively coupled plasma mass spectrometry (ICP-MS)

20 - 50 mg of dried material was digested in 1 ml of 67 % nitric acid overnight. Samples were heated to 95 °C until the solutions became clear. After cooling, the samples were centrifuged at 4,000 rpm for 30 minutes at 4 °C. 600 µl of the clear supernatant were combined with 7.44 ml of ddH2O and the solution was stored at 4 °C. ICP-MS analysis (Agilent 7700, Agilent Technologies, Waldbronn, Germany) was conducted at the metabolomics platform, Biocentre of the University of Cologne, relating elemental contents to dry weight. Total plant mineral contents were calculated using a root-to-shoot ratio of 1:2 (O’Sullivan et al. 2022, Tiziani et al. 2020).

### Quantification of phosphate

Inorganic and total phosphate were quantified according to Chiou et al. (2006). 5 mg of lyophilized plant powder was dissolved in 250 µl extraction buffer (10 mM Tris, 1 mM EDTA, 100 mM NaCl, 1 mM 2-mercaptoethanol, pH: 8.0). 100 µl of extract was mixed with 900 µl of 1 % glacial acetic acid and incubated at 42 °C for 30 min. 300 µl of this mixture was used for the quantification of inorganic phosphate. To 100 µl of this mixture, 30 µl of 10 % Mg(NO3)2 was added and then incinerated to ash. After cooling the ash was dissolved in 300 µl of 0.5 N HCl and total phosphate was quantified. To quantify phosphate, 700 µl assay solution (0.35 % NH4MoO4, 0.86 N H2SO4, and 1.4 % ascorbic acid) was added to 300 µl of phosphate extract and incubated at 42 °C for 30 min. The phosphate content was quantified at 820 nm using a plate reader.

### Quantification of photosynthetic parameters

FlourPen (FP110-LM/X, Photon System Instruments, Drásov, Czech Republic) protocols NPQ2 and LC3 were used for pulse amplitude modulation fluorometry. Before each measurement, cotyledons or leaves of 12-day-old plants were dark adapted by covering with leaf clips for 15 min.

Photosynthesis and respiration rates were measured under ambient conditions using an Oxygraph+ with LeafLab2 (Hansatech Instruments Ltd, Norfolk, UK). 200 µl of 1 M NaHCO3 was added to each measurement chamber. Samples were light adapted to 200 µmol photons · s^-1^ · m^-2^ for 7 min and respiration rates were measured in 500 and 1000 µmol photons · s^-1^ · m^-2^ light were measured followed by respiration rate measurement in the dark.

Leaf area and dry weight were measured.

### Protein extraction, digestion and sample preparation for proteome analysis via mass spectrometry

The procedure is based on the single-pot solid-phase-enhanced sample preparations (SP3) protocol for proteomics experiments of Hughes et al. (2019) and Mikulášek et al. (2021). The detailed adapted protocol, which was used here, can be found in Angermann et al. (2024). In brief, proteins were solubilized with sodium dodecyl sulfate (SDS) from lyophilized plant powder. Disulfide bridges were reduced with dithiothreitol (DTT) and subsequently alkylated with iodoacetamide (IAM). Equal shares (v/v) of hydrophobic and hydrophilic magnetic beads (No. 441521050250, No. 241521050250, Sera-Mag, Cytiva, Marlborough, Massachusetts, USA) were mixed to bind the denatured proteins. The beads were washed multiple times with 80 % ethanol to get rid of interfering components for the digestion. Finally, the proteins were digested overnight with trypsin (Mass Spectrometry Grade, Promega Corporation, Madison, Wisconsin, USA). The peptide-containing supernatants were collected and desalted on 50 mg Sep-Pak tC18 columns (Waters, Milford, Massachusetts, USA). The purified peptides were quantified with a quantitative Peptide Assay Kit (Thermo Fisher Scientific, Rockford, Illinois, USA) and adjusted to 400 ng · µl^-1^ in 0.1 % formic acid.

### Quantitative Shotgun Proteomics by Ion Mobility Mass Spectrometry (IMS-MS/MS)

For the cotyledon timeline experiment and the cotyledon removal experiment 400 ng peptides were injected via a nanoElute1 UHPLC (Bruker Daltonics GmbH, Bremen, Germany) and separated on an analytical reversed-phase C18 column (Aurora Ultimate 25 cm x 75 µm, 1.6 µm, 120 Å; IonOpticks, Collingwood, Victoria, Australia). Using MS grade water and a multi-staged gradient acetonitrile containing 0.1 % formic acid (0 min, 2 %; 54 min, 25 %; 60 min, 37 %; 62 min, 95 %; 70 min, 95 %), peptides were eluted and ionized by electrospray ionization with a CaptiveSpray1 ion source and a flow rate of 300 nl · min^-1^. The ionized peptides were separated, fragmented and analyzed with a standard data-dependent acquisition parallel accumulation–serial fragmentation application method (DDA-PASEF) of the system with the following settings: Ion mobility window: 0.6 – 1.6 V · s · cm^-^², 10 PASEF ramps, target intensity of 20,000 (threshold 2,500), and a cycle time of ∼1.1 s. The analysis was performed on a timsTOF-Pro2 mass spectrometer (Bruker Daltonics GmbH, Bremen, Germany).

For the comparison of cotyledons with true leaves (Fig. 3, Supplementary dataset S3) 400 ng of peptides were injected via a nanoElute2 UHPLC (Bruker Daltonics GmbH, Bremen, Germany) and separated on the same type of analytical column as described above. Using the same multi-staged gradient as above, but here peptides were ionized by a CaptiveSpray2 ion source and analyzed on a timsTOF-HT mass spectrometer (Bruker Daltonics GmbH, Bremen, Germany), which had following DDA-PASEF settings: Ion mobility window of 0.7 – 1.5 V · s · cm^-^², 4 PASEF ramps, target intensity 14,500 (threshold 1,200), and a cycle time of ∼0.53 s.

### Data Processing and Functional Annotation

The ion mobility spectrometry (IMS)–MS/MS spectra from all experiments were analyzed with the MaxQuant software (Cox and Mann 2008) using default search parameters and the proteome database of *Lupinus albus* published by Xu et al. (2020) on UniProt.org (UP000464885) for protein identification. The calculation of label-free quantification (LFQ) values and intensity-based absolute quantification (iBAQ) values were both enabled. Data evaluation was performed using Perseus (Tyanova et al. 2016). Proteins were excluded from further analysis if they were not detected in at least four out of five biological replicates in at least one of the sample groups. Subsequently, missing values were replaced with randomly chosen low values from a normal distribution. Significant changes were calculated from the LFQ values using Student’s t-tests (p = 0.05). Furthermore, a fisher exact tests were performed to identify significantly enriched or depleted metabolic pathways. A recently published annotation database of *Lupinus albus* was used to obtain further information such as subcellular localization and metabolic pathway involvement (Angermann et al. 2024).

### Statistical analysis

Statistical analysis was performed using Student’s t-tests (two-tailed test, pooled variance). Sample groups were compared with the control (cotyledons: day 0; remaining plants: day 28) at p = 0.05.

### Author contributions

TMH, CA, and BH designed the research; CA and BH performed and evaluated the shotgun proteomics experiments; CA and BH measured and evaluated amino acid profiles; BBN quantified phosphate; HJM conducted experiments and analyzed data for mineral nutrient quantification; CA performed all the other experiments; TMH, CA and BH analyzed the data; TMH and CA wrote the paper with support from all other authors; HJM and PB wrote and corrected text; TMH agrees to serve as the author responsible for contact and ensures communication.

## Data availability

The mass spectrometry proteomics data have been deposited to the ProteomeXchange Consortium (http://proteomecentral.proteomexchange.org) via the PRIDE partner repository (Perez-Riverol et al. 2022) with the dataset identifier PXD054416.

## Supplementary Data

Supplementary Figure S1: Scheme of the experimental set-up.

Supplementary Figure S2: Protein abundance profiles of functional protein categories during *Lupinus albus* cotyledon development.

Supplementary Figure S3: Free amino acid contents of cotyledons and true leaves at day 12.

Supplementary Figure S4; Mineral contents of shoots and roots at day 28.

Supplementary Figure S5; Nitrogen saving strategies after removal of the cotyledons.

Supplementary Dataset S1: Metabolite profiles of cotyledons and 28-day-old plants.

Supplementary Dataset S2: Reorganization of the cotyledon proteome during transformation from storage organ to photosynthetic tissue: cotyledon proteome data.

Supplementary Dataset S3: Metabolite profiles and proteome data of true leaves vs. cotyledons at day 12.

Supplementary Dataset S4: Plant proteome at day 28 after detaching the cotyledons at different timepoints.

## Acknowledgments

We thank Pascal Klein for preliminary work on this project, Christina Mack and Dagmar Lewejohann for skillful technical assistance and Tim Hildebrandt for being curious and asking the right questions. We thank Dr. Alberto Andrino de la Fuente for support during C/N analysis. The authors thank the MS Platform and Sabine Metzger, University of Cologne, for conducting and providing support in the ICP-MS measurements. We thank Lorna Egan-Andreou for the graphic design of the lupin development scheme.

