## Supplementary Figures S1-5 for "Balancing nutrient remobilization and photosynthesis: the dual role of lupin cotyledons after germination"

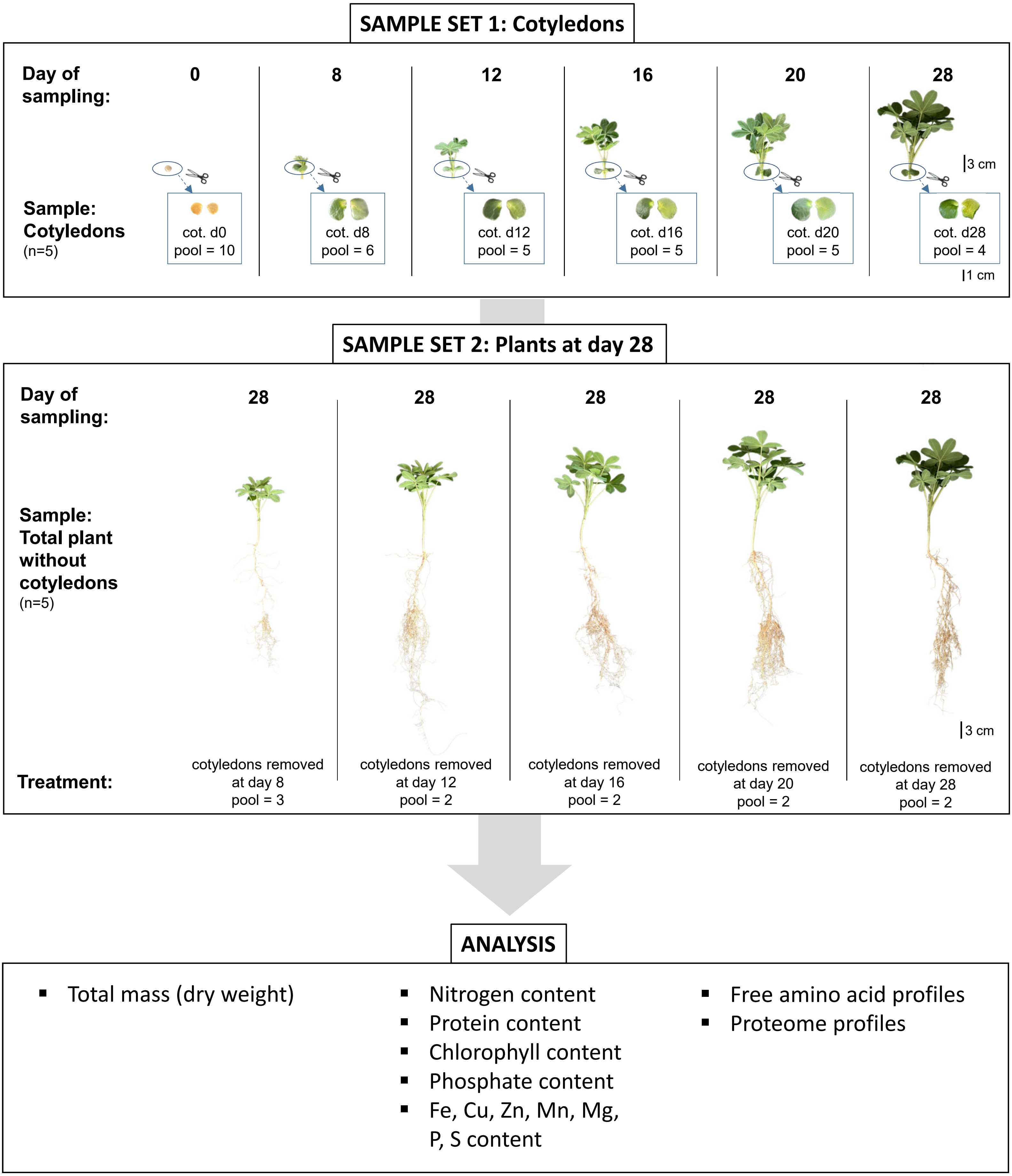


**Supplementary Figure S1.** Scheme of the experimental set-up: *Lupinus albus* plants were grown for twenty-eight days the absence of rhizobia and without nitrogen fertilization. The cotyledons were removed from subsets of these plants at day 8, 12, 16, 20 or 28 after sowing and harvested for subsequent analysis. Cotyledons of seeds were harvested as control. The remaining plants without cotyledons were harvested for subsequent analysis at day 28. The complete dataset is provided as Supplementary datasets S1 and S2.

**A**

| Functional protein categories | Number of proteins | Protein [mg · g $DW^{-1}$] | | | | | |
|---|---|---|---|---|---|---|---|
| | | 0d | 8d | 12d | 16d | 20d | 28d |
| Storage proteins | 23 | 93.7 | 45.7 | 25.4 | 12.8 | 9.5 | 11.4 |
| Protein metabolism | 1005 | 47.0 | 42.1 | 40.5 | 36.2 | 34.0 | 31.3 |
| Lipid metabolism | 220 | 46.9 | 19.5 | 16.4 | 11.8 | 9.7 | 8.9 |
| Vesicle trafficking | 134 | 1.6 | 1.2 | 1.4 | 1.3 | 1.1 | 1.2 |
| Cell | 323 | 33.7 | 12.2 | 11.4 | 9.2 | 8.0 | 8.6 |
| Stress | 169 | 15.5 | 9.3 | 9.1 | 7.7 | 7.9 | 8.9 |
| Redox | 131 | 11.2 | 8.9 | 9.1 | 8.1 | 7.6 | 7.9 |
| Secondary metabolism | 115 | 2.5 | 2.4 | 2.4 | 2.4 | 2.1 | 1.8 |
| Amino acid metabolism | 167 | 2.3 | 5.3 | 5.4 | 5.1 | 4.7 | 4.1 |
| Carbohydrate metabolism | 270 | 11.9 | 12.7 | 13.4 | 12.6 | 11.3 | 10.8 |
| Cellular respiration | 200 | 9.5 | 16.2 | 18.0 | 16.6 | 15.1 | 14.3 |
| Coenzyme metabolism | 111 | 1.0 | 2.6 | 2.2 | 2.0 | 1.8 | 1.8 |
| Nucleotide metabolism | 73 | 1.2 | 1.6 | 1.8 | 1.7 | 1.7 | 1.7 |
| Photosynthesis | 191 | 3.1 | 45.1 | 58.9 | 62.9 | 62.3 | 62.2 |
| Signalling | 79 | 0.9 | 1.2 | 1.5 | 1.5 | 1.5 | 1.8 |
| DNA | 64 | 2.0 | 2.2 | 2.9 | 3.3 | 3.8 | 4.2 |
| RNA | 313 | 2.8 | 3.1 | 3.3 | 3.2 | 3.3 | 3.2 |
| Nutrient uptake | 19 | 0.2 | 0.3 | 0.3 | 0.2 | 0.3 | 0.4 |
| Hormone metabolism | 73 | 1.6 | 1.2 | 1.6 | 1.5 | 1.5 | 2.0 |
| Transport | 180 | 7.0 | 6.7 | 6.8 | 6.7 | 6.6 | 7.1 |

**B**

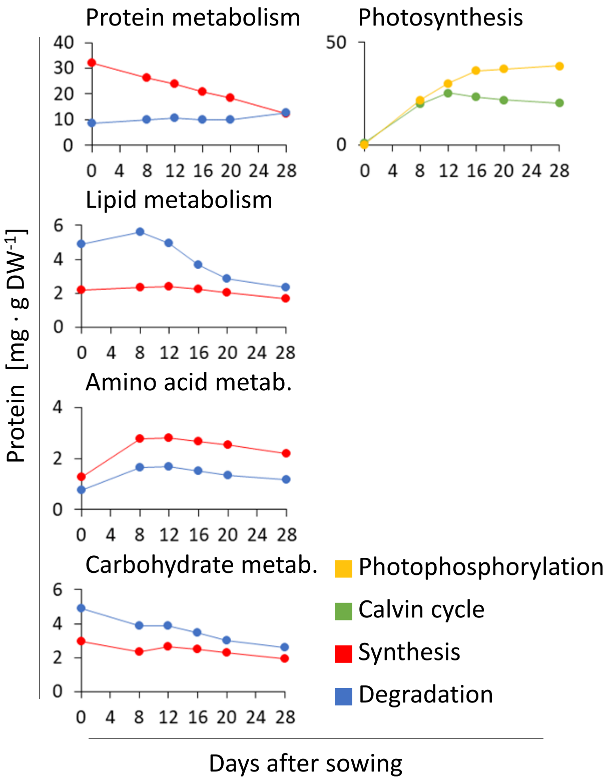


**Supplementary Figure S2:** Protein abundance profiles of functional protein categories during *Lupinus albus* cotyledon development. (**A**) Relative protein abundance of functional protein categories and number of different proteins detected in the respective category. Categories were adapted from MapMan4. (**B**) Protein abundance profiles of selected functional categories. The complete dataset is provided as Supplementary dataset S2.

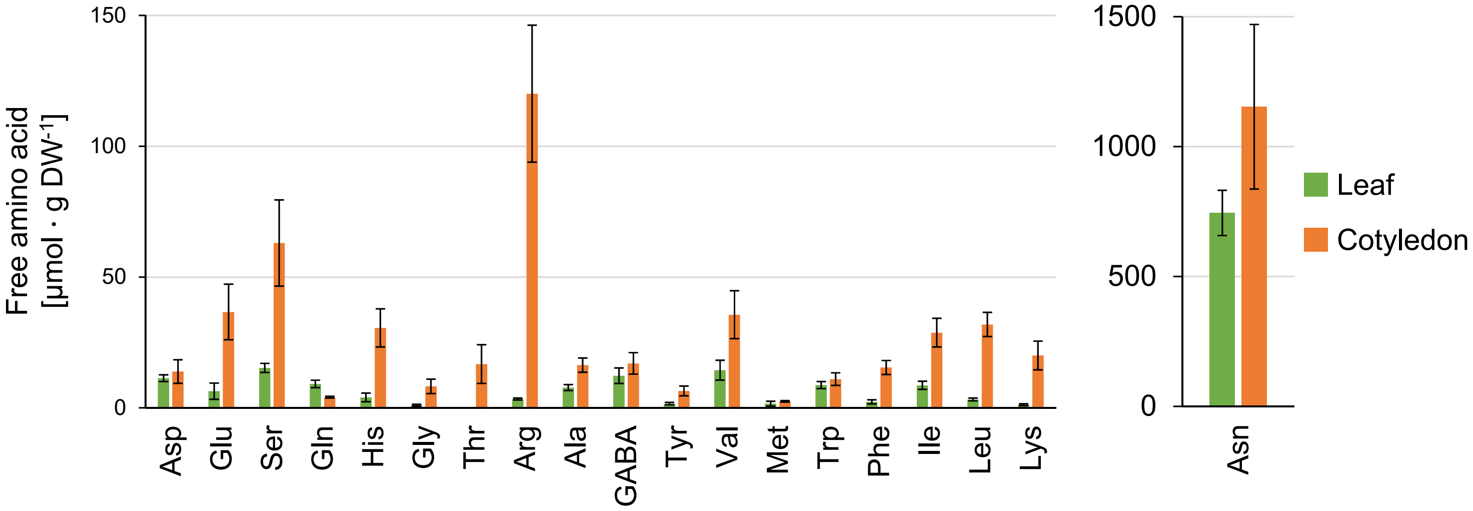


**Supplementary Figure S3:** Free amino acid contents of cotyledons and true leaves at day 12. The complete dataset is provided as Supplementary dataset S3.

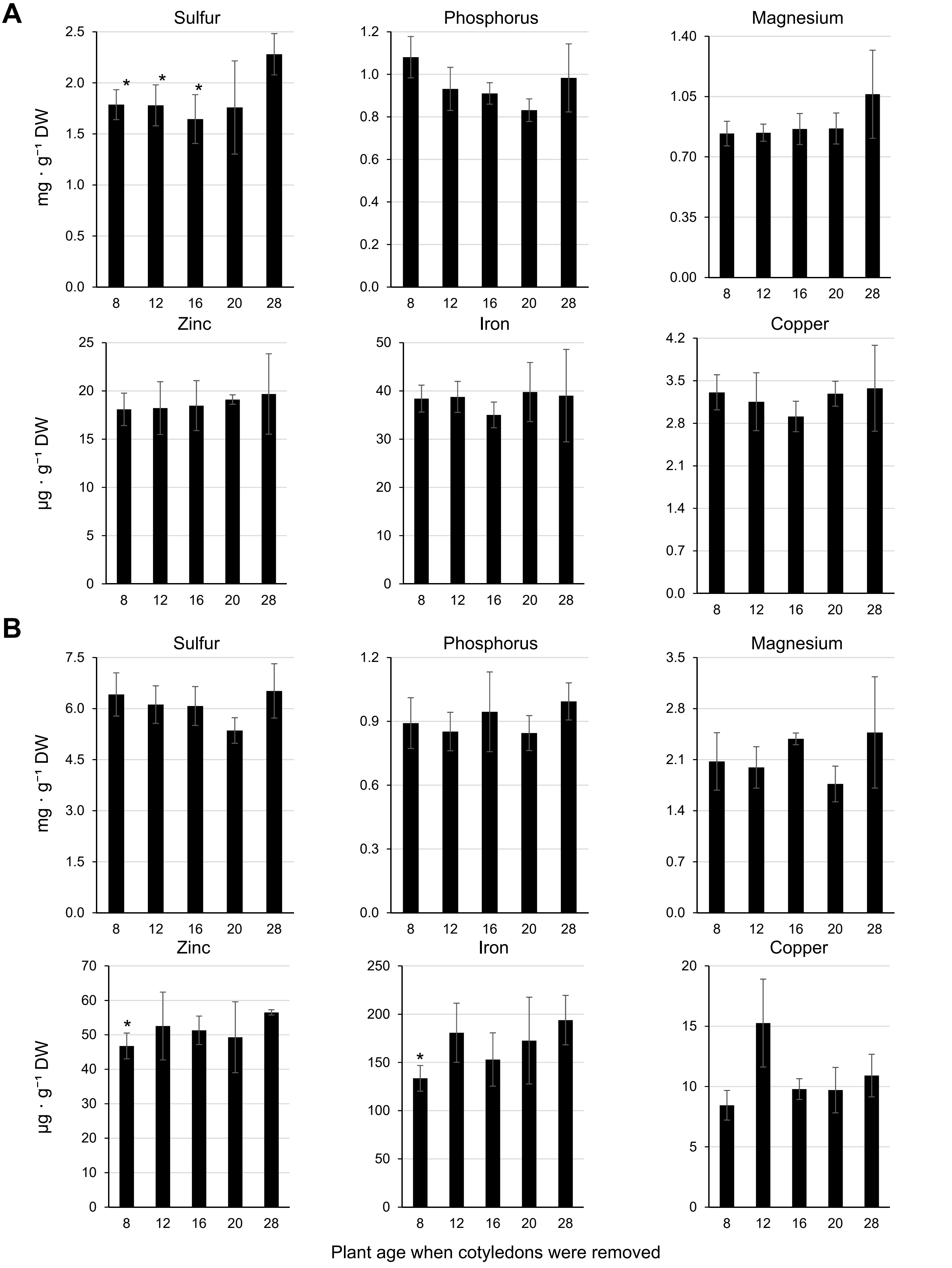


**Supplementary Figure S4:** Mineral contents of shoots and roots at day 28. (**A**) Shoot. (**B**) Root. Data presented are means ± SD (n = 3). *significantly different to control (28d) ($p<0.05$). Complete dataset is provided as Supplementary dataset S1.

### Nitrogen saving strategies after removal of cotyledons:

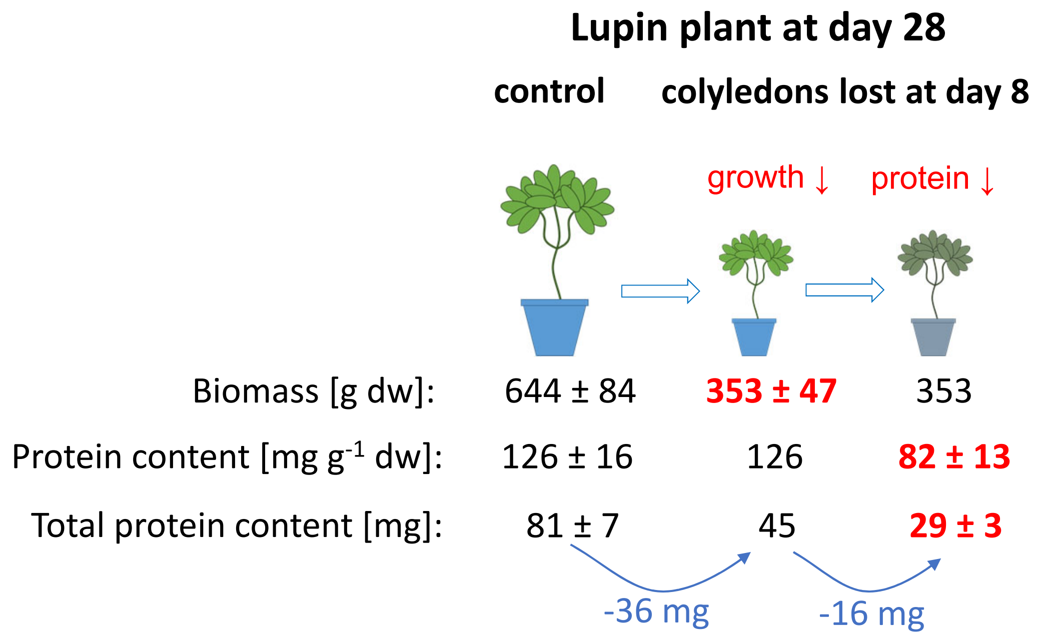


1. Reducing **plant biomass** from 644 to 353 mg dry weight

   ⇨ saves **36 mg protein**

2. Reducing **protein content** from 126 to 82 mg protein $g^{-1}$ dry weight

   ⇨ saves **16 mg protein**

**Supplementary Figure S5**: Nitrogen saving strategies after removal of the cotyledons: The plant is able to save 15.5 mg protein by reducing its relative protein content by 35 % from 126 ±16 to 82 ± 13 mg · g-1 DW after premature cotyledon loss at day 8. The most effective nitrogen saving strategy, however, is the reduction in plant biomass from 644 ± 84 to 353 ± 47 mg dry weight that, based on the mean tissue protein content of the control plant, would save 36.7 mg protein.
